# Assemblies of F-actin and myosin-II minifilaments: steric hindrance and stratification at the membrane cortex

**DOI:** 10.1101/656082

**Authors:** Amit Das, Abrar Bhat, Rastko Sknepnek, Darius Koster, Satyajit Mayor, Madan Rao

## Abstract

Recent *in-vivo* studies have revealed that several membrane proteins are driven to form nanoclusters by active contractile flows arising from F-actin and myosin at the cortex. The mechanism of clustering was shown to be arising from the dynamic patterning of transient contractile platforms (asters) generated by actin and myosin. Myosin-II, which assemble as minifilaments consisting of tens of myosin heads, are rather bulky structures and hence a concern could be that steric considerations might obstruct the emergence of nanoclustering. Here, using coarse-grained, agent-based simulations that respect the size of constituents, we find that in the presence of steric hindrance, the patterns exhibited by actomyosin in two dimensions, do not resemble the steady state patterns observed in our *in-vitro* reconstitution of actomyosin on a supported bilayer. We then perform simulations in a thin rectangular slab, allowing the separation of a layer of actin filaments from those of myosin-II minifilaments. This recapitulates the observed features of *in-vitro* patterning. Using super resolution microscopy, we find direct evidence for stratification in our *in-vitro* system. Our study suggests the possibility that *molecular stratification* may be an important organising feature of the cortical cytoskeleton *in-vivo*.

## I. INTRODUCTION

The transient nanoclustering of many cell surface molecules is strongly influenced by interaction with the actomyosin cortex [1–7], a thin layer of actin cytoskeleton and myosin motors (non-muscle myosin-II) [8] that is juxtaposed with the membrane bilayer. While the thickness of the cortical actin layer has been measured to be around 250 nm (in HeLa cells [9]), its ultrastructure is as yet poorly defined, although evidence suggests that it comprises both dynamic filaments [3] and an extensively branched static meshwork [10]. As a consequence, the action of myosin motors on the actin filaments at the cortex [11] drives the local clustering of cell surface proteins that bind to it [1–3].

This layered structure of a multicomponent, asymmetric bilayer juxtaposed with a thin cortical actomyosin layer forms the basis for a model of the cell surface as an *Active Composite* [1, 3, 5]. In these earlier studies [3, 12], we had used a coarse grained description, based on active hydrodynamics [13], that is agnostic to molecular details. Recently, it has been shown that the statistics and dynamics of clustering is recapitulated in a minimal *in vitro* reconstitution of a supported bilayer in contact with a thin layer of short actin filaments and Myosin-II minifilaments (both of skeletal muscle origin), driven by the hydrolysis of ATP [14, 15]. The success of this minimal setup motivates us to revisit our continuum hydrodynamic description from a more molecular, albeit coarse-grained, standpoint.

Non-muscle Myosin-II, which also assemble as minifilaments [16–19] and consists of around 30 myosin heads, are rather bulky structures [14], and hence a concern could be that steric considerations might obstruct the emergence of membrane protein nanoclustering. To address this, we perform a coarse-grained, agent-based Brownian dynamics simulation of a mixture of polar active filaments built from “actin monomers” and Myosin-II minifilaments comprising multiple “myosin heads”. Our coarse-grained model respects the relative sizes of the individual constituents and hence incorporate crucial steric effects. We study the phase diagram characterising actomyosin patterns in a strictly two-dimensional (2d) setting, upon changing F-actin length, myosin minifilament concentration and the strength of the contractile force exerted by individual myosin-II motors on actin filaments. We compare the patterns that we obtain in simulations with the steady state patterns obtained in an *in-vitro* reconstitution of a thin layer of actomyosin on a supported bilayer, first described in Ref. [14]. We find that steric effects are indeed significant and the patterns obtained do not resemble the steady state patterns in the *in-vitro* reconstitution system. We then perform simulations in a thin rectangular slab, allowing the separation of a layer of actin filaments from those of myosin-II minifilaments. This recapitulates the *in-vitro* patterning - using super resolution microscopy, we look for and find direct evidence for stratification in the *in-vitro* system.

## II. MODEL

### A. Agent-based simulations of F-actin and myosin-II minifilaments

We model both F-actin and myosin-II minifilaments [17–19] as coarse-grained semi-flexible polymers using a bead-spring representation with steric repulsion, Fig. 1a. The actin filament is a linear polymer, while the myosin-II minifilament is a polymer with side chains depicting a collection of myosin head domains attached to a backbone made up of tail domains [18]. We assign the same diameter for all the beads *σ* for simplicity, irrespective of whether it is a subunit of F-actin or myosin minifilament. Consistency with previously reported dimensions [14, 17, 20–23], implies that 1 F-actin bead corresponds to ≈ 40 G-actin monomers and 1 myosinhead bead corresponds to ≈ 3 − 4 Myosin-II heads. The monomers are connected by harmonic bonds. For convenience, we use the same stretch spring constant for all bonds. In addition, changing the angle between two consecutive bonds is penalised, effectively introducing a bending stiffness. We choose appropriate values for these elastic constants to ensure that the actin filaments have a persistence length larger than 16 *μ*m [24] and the myosin minifilaments have flexible heads attached to a very rigid backbone [25]. We keep track of the polarity of F-actin by labelling the terminal beads in each filament by a plus or barbed end (+) and minus end (-). The dynamics of the beads is described by overdamped, Brownian dynamics equations (see, *Supplementary Information (SI)* for details).

**FIG. 1.**
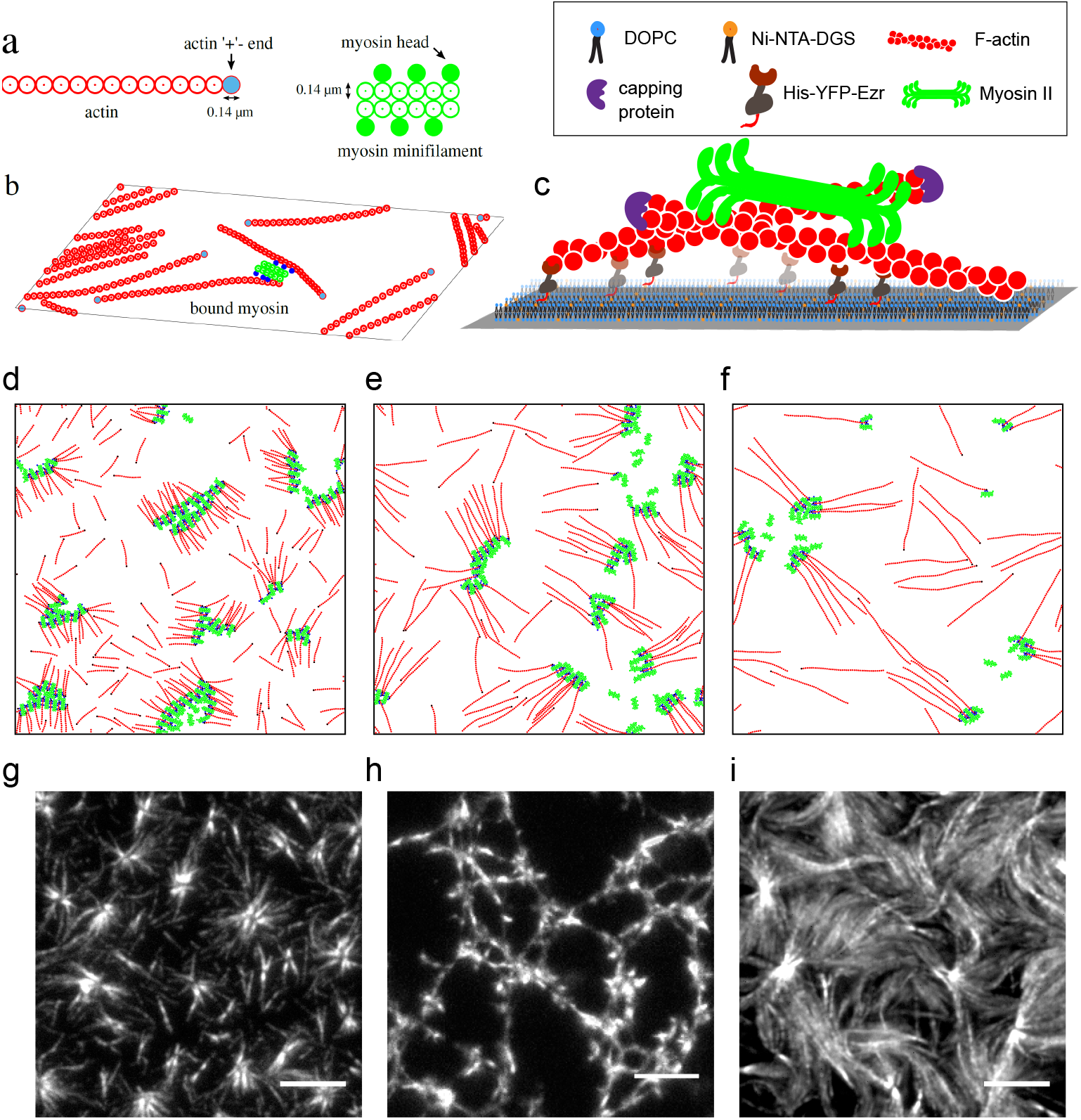
Ingredients of agent-based simulations in 2d and *in-vitro* reconstitution set up. (a) Schematic of actin filaments (red) and myosin minifilaments (green) used in the agent-based model, with real space dimensions indicated. One F-actin bead corresponds to 40 G-actin monomers and one myosin bead corresponds to ~ 3-4 heads. (b) Schematic showing a collection of F-actin and myosin minifilaments in 2d, with myosin motors bound to two actin filaments (bound myosin heads are coloured blue). (c) Schematic of the *in-vitro* reconstitution system showing the hierarchical assembly of a supported lipid bilayer, linker protein (HYE), capped actin filaments and muscle myosin II. (d-f) Typical simulation snapshots of different aster configurations observed in the strictly 2d-simulations, as a function of increasing F-actin length *l*_*a*_: (d) *l*_*a*_ = 2.32 *μ*m, (e) *l*_*a*_ = 4.84 *μ*m and (f) *l*_*a*_ = 7.19 *μ*m. The plus-ends of the actin filaments are coloured black. (g-i) Typical aster configurations observed in *in-vitro* experiments: isolated asters, connected asters and aster bundles, as a function of increasing F-actin length: (g) *l*_*a*_ = 2 *±* 1 *μ*m, (h) *l*_*a*_ = 3 *±* 1.5 *μ*m, and (i) *l*_*a*_ = 8 *±* 3 *μ*m. Scale bar = 5 *μ*m.

The myosin-head beads experience an attractive interaction with the F-actin beads, modelled by a Morse potential where the strength of a myosin-actin bond corresponds to the typical change in free energy associated with hydrolysis of one molecule of ATP [26]. In addition, each motor-head binds to an actin filament without any geometric restriction, applies an active force *f*_*a*_ on actin to displace itself in direction of the (+)-end of actin by sliding the actin in the opposite direction, and eventually unbinds. The binding/unbinding of each motor follows a simple Poisson kinetics [22]. Note that we allow a coarse-grained myosin bead to interact with many actin filaments. In addition to this, we allow for turnover of full myosin minifilaments, implemented by allowing the completely detached myosin minifilaments to escape from the medium and to reappear at a different location at a rate *k*_*m*_, while keeping the mean number of bound myosins fixed.

Our control parameters in the simulation are: (i) length of actin filaments *l*_*a*_, which we vary in the range of 2 − 9 *μ*m. (we have kept the length of the myosin-II minifilament fixed at ≈ 1 *μ*m), (ii) magnitude of active force *f*_*a*_ (related to the number of phosphorylated heads) for which we use values in range 1 − 9 pN per myosin head bead [20–23] and (iii) concentrations of myosin-II (*c*_*m*_) and F-actin (*c*_*a*_). We have introduced separate unbinding and binding rates *k*_*u*_ and *k*_*b*_, respectively, for the myosin head beads [22]. All other simulation parameters are held fixed across simulation runs. For other relevant details, see the *Methods* section and for a full description of the simulation model, the relevant parameters and their values, see the *SI*.

### B. In-vitro reconstitution experiments

The *in-vitro* reconstitution system consists of a thin actomyosin layer atop a supported lipid bilayer, as described in our earlier work [14]. Briefly, we prepare functionalised lipid membranes, followed by hierarchical addition of actin-binding linker protein (HYE), pre-polymerised actin filaments together with control of actin length by capping proteins, myosin II motors, and limited ATP (100 *μ*M) Fig. 1c. Actin filaments undergo spatial reorganisation in the presence of ATP-fuelled myosin activity, and reach a static steady state as a consequence of steady ATP depletion. We record these steady state patterns (asters) using total internal reflection fluorescence microscopy (see *Methods*). We have systematically tuned the actin filament density, actin filament length, and myosin density, and have recorded three distinct actomyosin patterns: isolated asters, connected asters and aster bundles, Fig. 1g-i.

## III. RESULTS

### A. Simulations in a strictly two-dimensional layer

We start with the agent-based simulations in a strictly two-dimensional actomyosin system, i.e., when both Factin filaments and Myosin-II motors reside in the same plane. As expected, steady state configurations displayed by the rigid F-actin filaments alone (with no myosin) fall into three broad classes, determined by an interplay between the filament length *l*_*a*_ and the filament concentration *c*_*a*_ [27–29], (i) dilute – when the rods are far apart, 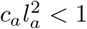 (ii) semi-dilute – when the rods entangle, 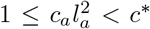 (the Onsager concentration for isotropic-nematic transition [30]); and (iii) concentrated – when the rods start to align, *c > c\**. In this work, we restrict ourselves to the dilute and semi-dilute regimes;

The initial configurations for the Brownian dynamics simulation of the full actomyosin system, are chosen from an ensemble where the actin and myosin filaments are uniformly distributed in position and orientation, at a fixed concentration of filaments. Typically, after a time *t* ≈ 15 − 30 s, as a result of the binding, active pushing and unbinding of myosin beads, the actin filaments and myosin are driven to form clusters, consistent with our earlier theoretical studies [3, 12], as well as simulations [20, 22, 31, 32], and *in-vitro* experiments [14, 33–39].

Figure 1d- 1f show typical configurations of clusters observed in simulations, as one increases the length of actin filaments. In comparison, we note that the clusters formed in the *in-vitro* reconstitution experiment (Fig. 1g-1i), go from being isolated asters, to connected asters to bundled asters, as the length of actin filaments is increased from 2 to 9 *μ*m, as reported earlier [14]. However, significant differences between the configurations observed in the strict 2d simulations and *in-vitro* experiments appear, when we focus on molecular patterning in the isolated clusters (Fig. 2).

**FIG. 2.**
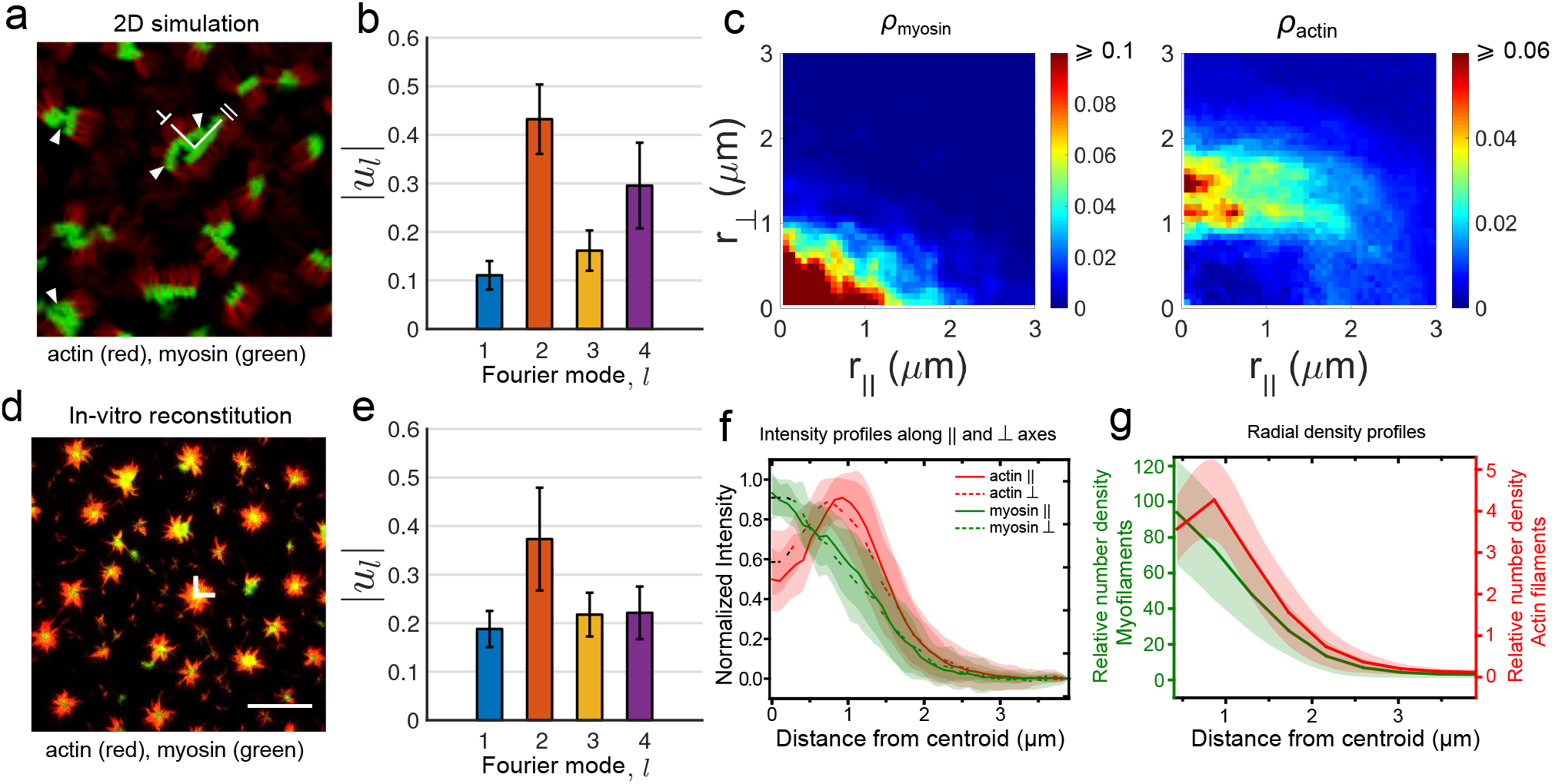
Comparison of cluster patterns in the strictly 2d simulations and *in-vitro* experiments. (a) Patterning of actomyosin clusters in the dilute limit obtained from agent-based simulations in the strictly two-dimensional geometry. These patterns have been obtained by coarse-graining the density of actin (red) and myosin (green) on a 2d grid with spacing 2 *σ*. The relevant parameters are: *l*_*a*_ = 2.32 *μ*m, *c*_*a*_ = 1.2 nM, *c_m_/c_a_* = 0.4, *f*_*a*_ = 2 pN and *k_u_/k_b_* = 0.1. Note that there are no overlapping regions (yellow) as a result of strong steric effects. (b) The shapes of most of the clusters observed in the simulations are strongly anisotropic, almost rectangular and some even with appearence of bi-stripes (marked by white arrows in panel a), as determined from a Fourier analyses of the shape (*Methods*). Note the large amplitude of the *l* = 2 and *l* = 4 modes compared to the *l* = 1 mode, averaged over *N* = 19718 clusters. (c) Density profiles of myosin and actin along the ⊥ and ‖ directions of the clusters also show strong anisotropy. Data taken over *N* = 24707 clusters. (d) Patterning of actomyosin clusters in the dilute aster limit obtained from the *in-vitro* experiments, with actin (red), myosin (green) and a distinct region of overlap (yellow). Scale bar = 10 *μ*m. (e) In contrast to the strictly 2d simulations, the shapes of most of the clusters are largely circular as shown by the relatively elevated amplitude of the *l* = 1 mode compared to the higher modes. Data averaged over *N* = 561 clusters. (f) Normalised intensity profile (*I/Imax*) extracted from TIRF images of actin and myosin along orthogonal directions (⊥, ‖) confirming isotropy of the asters. Data averaged over 20 asters, across 4 different experiments. (g) Circularly averaged radial density profiles of actin and myosin show enrichment of the latter at the core (calibration of TIRF images in Fig. S3). Data averaged over 49 asters, across 4 different experiments. Error bars denote standard deviation.

Most of the actomyosin clusters observed under these simulation conditions are anisotropic, many of them forming rectangular bi-stripes, with actin on one side and myosin on the other (Fig. 2a). On the other hand, the clusters in the *in-vitro* experiments, are isotropic or circular, with myosin at the core and actin filaments extending radially outward, with the (+)-end oriented towards the core (Fig. 2d, 2g). The difference between the strict 2d simulations and *in-vitro* experiments, is apparent in the Fourier spectrum of the cluster shape (Fig. 2b, 2e) and in the density profiles of myosin and actin (Fig. 2c, 2f). Interestingly, when we remove the steric interactions in our simulations, the resulting patterns begin to resemble the experimental patterns (Fig. S1). It is clear that the strong steric hindrance of the bulky myosin filaments gets in the way of generating the actomyosin patterns that are potentially responsible for cell surface clustering and bringing membrane proteins in close proximity [3, 14].

### B. Simulations in a stratified actomyosin layer

To circumvent the frustration arising from the unavoidable steric constraints, we perform simulations in a stratified actomyosin layer, with actin filaments distributed in the xy-plane at *z* = 0 and myosin minifilaments restricted within a rectangular slab of thickness *z*_*s*_ = 3 *σ*, by a harmonic potential *U* = 1/2 *k*_*c*_(*z* - *z*_0_)^2^ centred at *z*_0_ = *z*_*s*_/2.

We find that stratification indeed releases the frustration from steric constraints and the steady state patterns of actomyosin start resembling the patterns seen *in-vitro* (Fig. 3a-c). The clusters are now more circular, with myosin concentrated at the cores, closely resembling the *in-vitro* patterning (Fig. 3b, 3c). Encouraged by this, we study the change in steady state patterns as a function of actin and myosin concentrations and length of actin filaments. For this, we set *f*_*a*_ = 2 pN, *k*_*u*_/*k*_*b*_ = 0.2 and work with a large enough *k*_*b*_, the myosin binding rate on actin, so that the eventual remodelling of the aster configurations happens at timescales much larger than typical simulation runs. The resulting phase diagram is qualitatively similar to the one in Ref. [14].

**FIG. 3.**
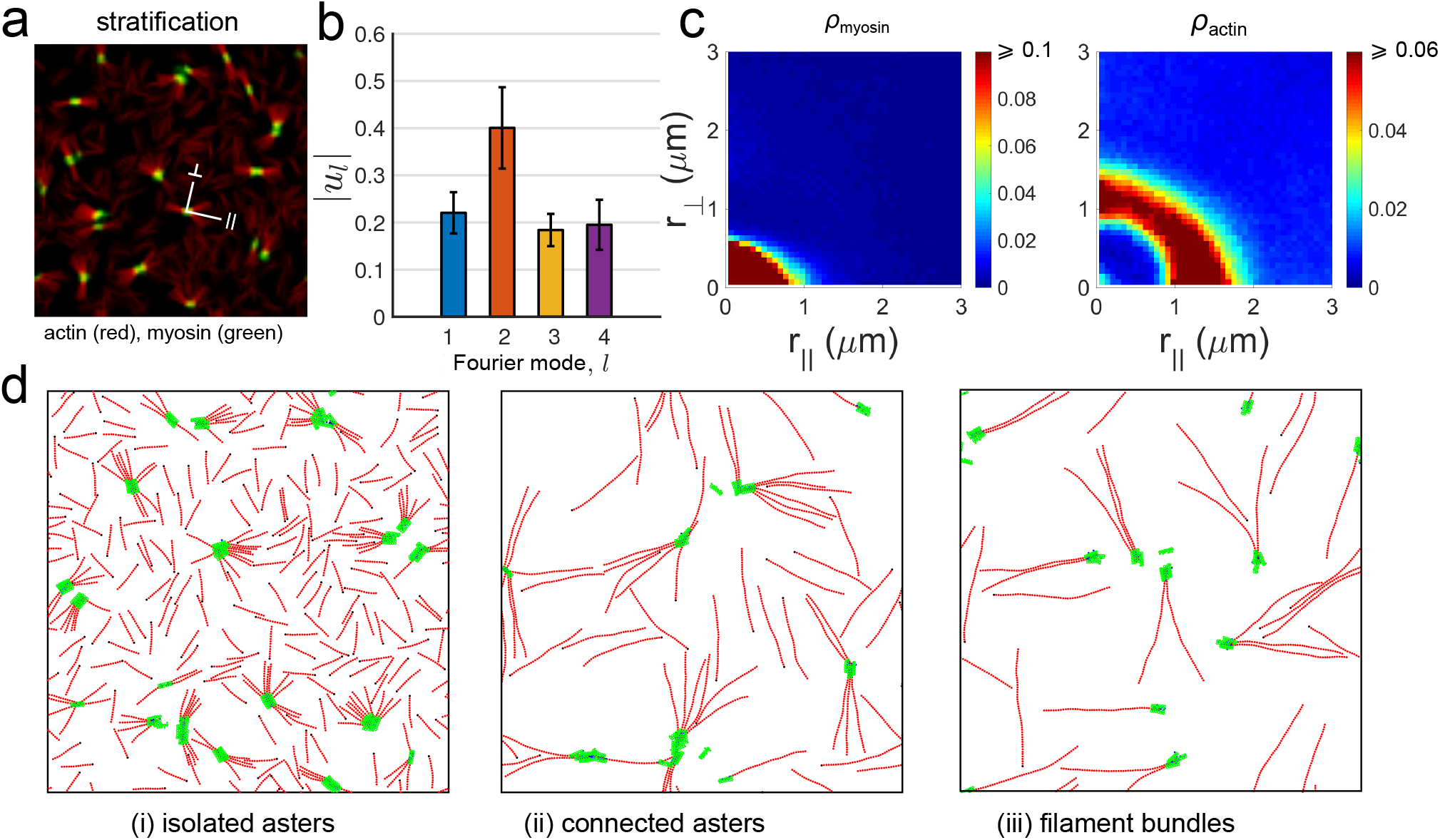
Simulations with stratification. (a) Patterning of actomyosin clusters in the dilute limit obtained from simulations in the stratified geometry. The parameters are *l*_*a*_ = 2.32 *μ*m, *c*_*a*_ = 1.5 nM, *c*_*m*_/*c*_*a*_ = 0.2, *f*_*a*_ = 2 pN and *k*_*u*_/*k*_*b*_ = 0.2. In this case, we see a co-localization of myosin and actin when projected in the x-y plane. (b) As in the *in-vitro* study, the clusters are more circular, as shown by the relatively elevated amplitude of the *l* = 1 mode compared to the higher modes. Data averaged over *N* = 33518 clusters. (c) Density of myosin and actin along the ⊥ and ‖ directions of the clusters show isotropic profiles, with myosin enriched at the cores and the *relative concentration* of actin enriched at the periphery. Data taken over *N* = 44537 clusters. (d) Representative snapshots of the different phases obtained on increasing actin filament length, (i) *l*_*a*_ = 2.32 *μ*m, (ii) *l*_*a*_ = 6.02 *μ*m, (iii) *l*_*a*_ = 8.37 *μ*m, showing isolated asters, connected asters and filaments bundles.

At the concentrations of actin under study, the steady state patterns are determined by a competition between the nematic alignment, parametrised by the Frank splay elastic constant *K*_1_ and a splay instability induced by myosin contractility [40]. The splay elastic constant depends linearly on the filament concentration and filament aspect ratio [41], thus when the filament concentration is low, and the filaments lengths are small, ≤ 6*μ*m, the splay instability due to myosin contractility dominates. As shown in Ref. [40], this leads to the formation of an *isolated aster* phase with the (+)-ends of actin filaments in each aster pointing inwards.

As one increases the length or concentration of actin filaments, the system transitions from an isolated aster phase in the dilute regime to a semi-dilute regime. To understand the structures that form in the semi-dilute regime, we first note that because of the active contractile stresses, there emerge two new length scales - the aster size and the aster separation ℓ* - both functions of the filament length, concentration and contractility. As we increase the actin filament length (or concentration), while keeping the other control parameters fixed, we obtain configurations where the long filaments span across the asters. Upon increasing myosin density, these spanning filaments via myosin, result in *connected asters* with altered polarity patterns, that we will discuss below.

Upon increasing the length or concentration of actin filaments further, the aligning tendency of the filaments starts to dominate over the contractility induced splay instability. This is because the splay elastic constant *K*_1_ depends linearly on the filament aspect ratio and filament concentration [41]. This leads to the formation of *filament bundles* of actin beyond *l*_*a*_ > 6 *μ*m, amidst a dilute dispersion of bundled asters. This is the active analogue of the entangled phase in passive mixtures of filaments and crosslinkers [42].

To display these changes in filament-motor configurations as a phase diagram, we need to arrive at a suitable order parameter. Since the various aster configurations are accompanied by filament-polarity sorting, an appropriate order parameter can be extracted from the behaviour of the pair-correlation function *g*_*m*−_(*r*) between the bound myosin heads and the (−)-end of actin filaments (Fig. 4a and Fig. S2) (See *Methods* section for definition). In the isolated aster phase, myosin heads are proximal to the (+)-ends of actin filaments, consequently the correlation between the myosin heads and (-)-end of the filaments will peak at a separation *r* = *l*_*a*_. In the connected aster phase, a fraction of the myosin heads that connect across asters are proximal to the (-)-end of actin; this will give rise to a new peak at at *r* = *σ*, the minimum separation of two beads. Finally, in the filament bundle phase, the peak height at *r* = *σ* relative to that at *r* = *l*_*a*_ decreases. Defining *R* ≡ *g*_*m*−_(*r* = *σ*), we record a non-monotonic variation as one sweeps across the three phases upon tuning *l*_*a*_ and *c*_*m*_/*c*_*a*_ (Fig. S2). This allows us plot the phase diagrams shown in Fig. 4b-c.

**FIG. 4.**
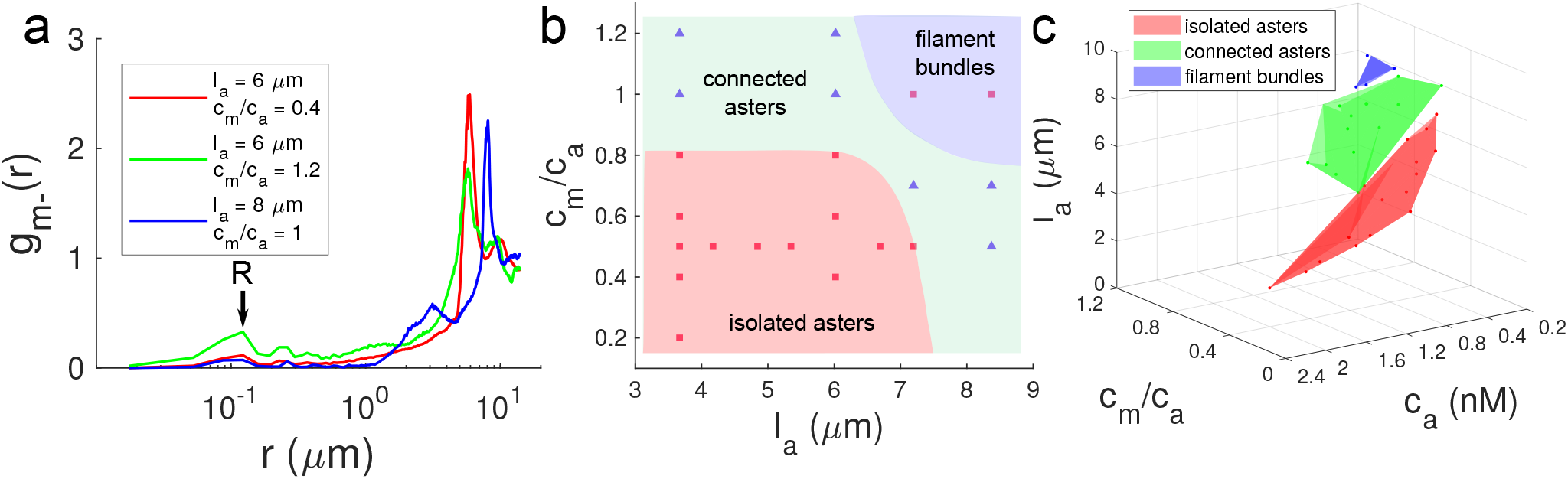
Characterization of the phases observed in our simulations of stratified actin filaments and myosin minifilaments. (a) Plots of *g*_*m*−_(*r*) for three different cases depicting three different phases observed. (b) A 2d phase diagram showing three different phases observed in our simulations on the *c*_*m*_/*c*_*a*_ vs. *l*_*a*_ plane. The squares correspond to cases where *R* ≤ *R*_*c*_ and triangles to cases with *R* > *R*_*c*_ where *R*_c_ is a suitable value (0.15) fixed in order to draw the phase boundaries. The bundle phase is confirmed by calculating the orientational alignment of the bound actin filaments around the myosin core, defined by *φ* (see *Methods*), which changes from zero in a disordered configuration to 1 in a fully aligned configuration. (c) Full phase diagram of the different phases in the space of length of actin (*l*_*a*_), concentration of actin (*c*_*a*_) and ratio of myosin to actin concentrations (*c*_*m*_/*c*_*a*_). The phase points have been assigned based on plots as in (b).

The phase diagram as a function of actin filament length and concentration of actin and myosin (Fig. 4c), bears a qualitative resemblance to the *in-vitro* phase diagram in Ref. [14]. This resemblance and the similarity of the actomyosin patterning in asters, suggests that stratification indeed releases the frustration imposed by steric constraints. We next look for evidence of such a release in the *in-vitro* experiments, especially at the cores of the asters observed in Fig. 2d.

### C. STED microscopy on *in-vitro* system reveals stratification

To obtain evidence of stratification in our *in-vitro* setup, we need to probe the configurations of actin and myosin transverse to the plane of the actomyosin layer. To this end, we perform 3d stimulated emission depletion (STED) microscopy on the *in-vitro* system, prepared with Star-635 labelled actin and Star-580 labelled myosin-II, with a resolution of 70 nm (100 nm) in the lateral xy plane, and 150 nm (180 nm) in the transverse z direction, for actin-635 (myosin II-580), see *SI*. Analysis of the actin and myosin-II signal along the z-axis shows distinct stratified organisation of actin and myosin.

We see that the periphery of the aster along the xy plane, is relatively enriched in actin filaments, with few myosins attached (Fig. 5a-d). On the other hand, the aster core is relatively enriched in myosin-II; the actin filaments appear to protrude into the z-direction, with myosin-II forming a cap over it. We note that the separation between the actin protruding column and myosin-II at the core, ranges from 150 nm to 800 nm with a mean at 393 nm (Fig. 5d-e).

**FIG. 5.**
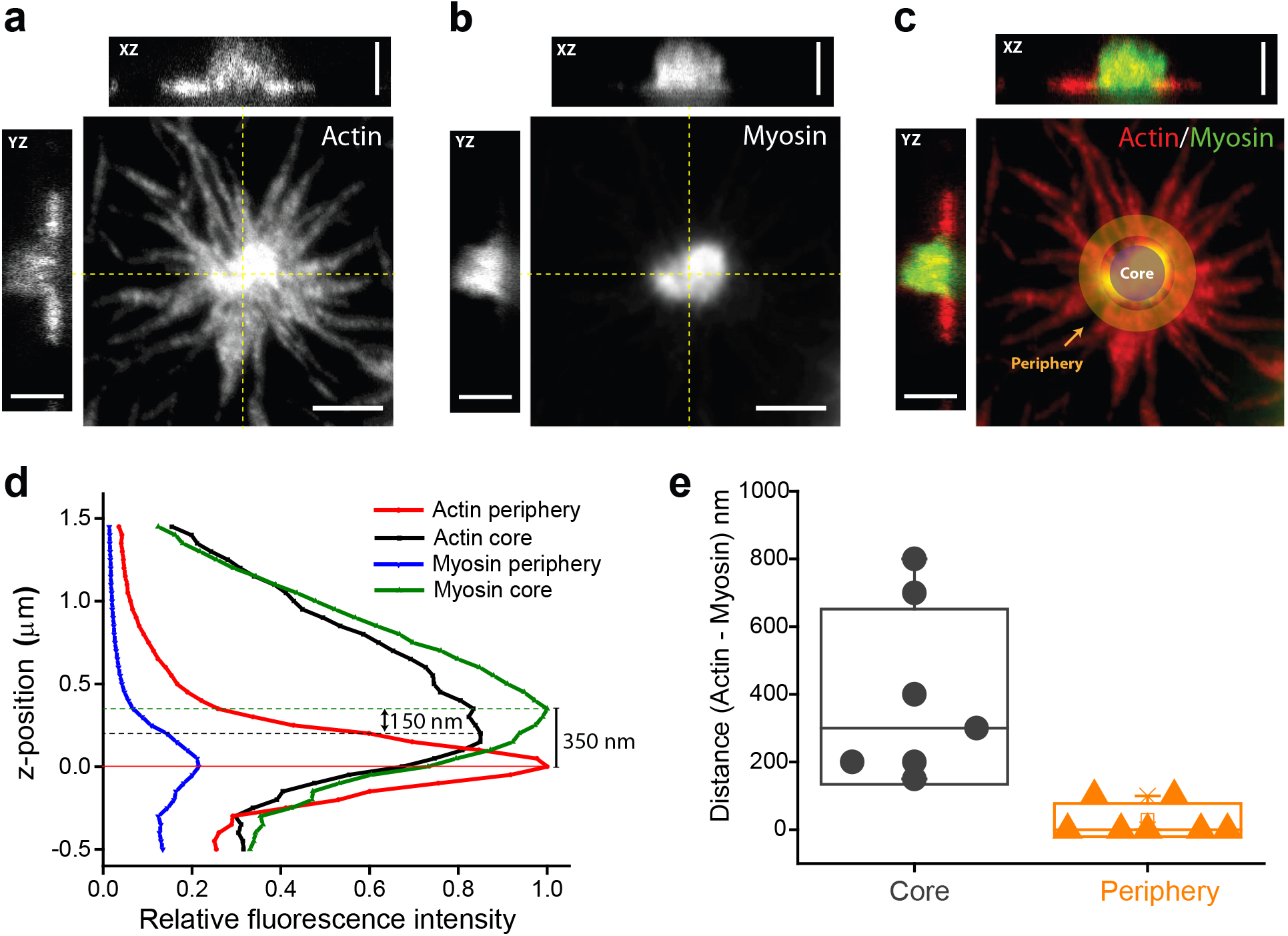
Stratified organisation of actin and myosin in asters. (a-c) 3d STED microscopy of an acto-myosin aster showing average intensity projection and XZ/YZ intensity profile of actin (a), myosin (b), and the merge (c) demarcating the core and the periphery used for subsequent measurements (Scale: *XY* = 2*μ*m; *XZ/Y Z* = 1.5*μ*m). (d) Intensity plot showing the relative intensities (*I/I*_*max*_) of actin and myosin along z-axis in the core (solid) and periphery (dashed). Actin intensity peak in the periphery was arbitrarily chosen as position 0. (e) Distance between actin and myosin intensity peaks (as shown in d) in the core and the periphery, quantified across 7 different asters.

Our observation suggests that the myosin enriched at the core, *draws in* the actin filaments, which protrude into the transverse direction (with respect to the bilayer). The results obtained from the STED measurements are an experimental confirmation of the stratification of the actomyosin layer.

## IV. DISCUSSION

We have presented results of detailed agent-based simulations of actomyosin assemblies that account for realistic steric restrictions, both in strict 2D and in stratified geometries. A key observation is that steric effects, often ignored in such simulations, can drastically affect the observed steady state configurations. We find that in order to recover the steady state configurations observed in *in-vitro* experiments [14], it is necessary to *stratify* the Factin and myosin-II minifilaments. We have constructed a phase diagram as a function of the concentrations of F-actin, myosin motors and the length of actin filaments in this actively driven system, which bears a satisfying resemblance to the steady state phase diagram reported in [14]. The generation of this phase diagram, resembling the *in-vitro* system, is a significant achievement of the simulations developed here. This reveals many of the key ingredients at the microscopic level and emergent mesoscale which recapitulate the *in-vitro* system.

Motivated by our agent-based simulations, we next look for evidence of stratification in an *in-vitro* reconstitution of actin and myosin on a supported bilayer. Using super resolution microscopy, we find direct evidence for stratification. We believe these observations have a bearing on the possible role of *molecular stratification* within the active cortex at the cell surface.

Molecular stratification and its regulation has been invoked as an organising principle in the elaboration of the focal adhesion complex [43–45] at the cell surface. In the context of the lateral nanoclustering of the outer cell surface GPI-anchored proteins, we demonstrated that the lipid-tethered protein, in conjunction with cholesterol, sphingolipids and inner leaflet phosphatidylserine (PS), forms a molecular bridge transverse to the asymmetric bilayer membrane, that helps connect the outer cell surface to the actomyosin cortex [46]. In this paper, we argue that a transverse organisation or stratification continues into the actomyosin cortex, thereby facilitating lateral nanoscale proximity of cell membrane proteins, that engage with the cortex. Our study shows that in the absence of stratification, steric effects would preclude such lateral nanoscale proximity. This has functional implications, since for example, cholesterol-dependent GPI-anchored protein nanoclustering is necessary for promoting integrin function [47, 48]. We take up some of these issues in a later study.

The molecular stratification that we observe is a consequence of steric effects. This is likely to be a key organizing element in many types of hierarchical self assembly and organization processes [43–45]. This naturally gives rise to stratification that is evident in many of the larger scale assemblies.

## ACKNOWLEDGMENTS

We thank Silke Henkes and Kees Weijer for many illuminating discussions. AD thanks School of Life Sciences, University of Dundee, UK for hospitality, and the Simons foundation for funding. MR and SM thank Jim Spudich for drawing attention to the role of steric interactions. We thank CIFF Imaging Facility at C-Camp, NCBS for the use of the STED microscope. RS thanks UK BBSRC (Award No. BB/N009789/1) for funding. SM and MR acknowledge a JC Bose Fellowship from DST (Government of India), and support from the NCBS-Max Planck Lipid Centre. SM acknowledges HFSP (RGP0027/2012), and Margadarshi fellowship (IA/M/15/1/502018) from The Wellcome Trust-DBT India Alliance.

## Methods

### Simulation details and units

In the agent-based model, we measure energy in units of *k*_*B*_*T* at the physiological temperature of 310 K, length in units of bead diameter *σ* ≈ 0.14 *μ*m and time in units of *τ* = *ζσ*^2^/*k*_*B*_*T* ≈ 10^−2^s where *ζ* is the friction coefficient of the medium associated with the drag on a monomer bead in the model [21]. Equations of motion are integrated using standard Euler-Maruyama algorithm with the time step *δt* = 1.5 × 10^−4^*τ*. The simulations are carried out up to 10^8^ time steps (≈ 10^4^*τ*) The simulations took 1 − 2 *×* 10^7^ steps to reach the steady state, which was identified by monitoring the autocorrelation functions of density and local orientation of the filaments. We perform simulations in 2d with all polymers restricted to the *xy*-plane, and also in a quasi-three-dimensional geometry with a stratified layer of myosin minifilaments placed “on top” of the plane of actin filaments. Dimensions of the 2d box in which all the actin filaments are restricted are: *L*_*x*_ = *L*_*y*_ = 28 *μ*m. Our simulations, done with the SAMoS program [51], include steric hindrance among all polymers via the repulsive Weeks-Chandler-Anderson (WCA) potential [52].

### Calculation of the order parameters for different phases

The pair-correlation function *g*_*m*−_(*r*) is a two-point density-density correlation function, defined as follows:

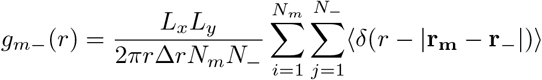

where *g*_*m*−_(*r*) represents the probability to find a myosin head, located at position **r**_*m*_ and an actin (−)-end, located at position **r**_−_, within a thin annular region of radius *r* and thickness ∆*r*. *N*_*m*_ and *N_−_* are the number of myosin heads and number of actin (−)-ends, respectively.

We calculate the local polar order parameter *φ* to quantify the orientational alignment of the bound actin filaments around a myosin core. We define *φ* to be the net orientation of actin filament segments (two consecutive actin beads bound by harmonic bond) found within 1 *μ*m of the centroids of bound myosin minifilaments. The formal definition of *φ* is the following [49] for *N*_*s*_ actin segments:

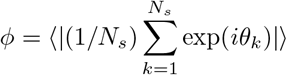

where the angular brackets represent time averages and *θ*_*k*_, the orientation of the *k*-th actin segment. In the isolated aster phase *φ* is small (≤0.5) because bound actins nearly form a closed loop of orientations around a bound myosin. However, *φ* slowly increases in the connected aster phase because of non-uniform populations of actin (+) and (−) ends around bound myosin. Finally, *φ* approaches unity in the phase dominated by actin bundles. This variation of *φ* has been used to confirm the phase boundary between the connected aster phase and the filament bundle phase of actin.

### In vitro reconstitution system

Our reconstitution system consisted of thin actomyosin layers coupled to supported lipid bilayers (SLBs) via linker proteins, as described in our earlier work [14]. Briefly, actin and myosin were purified from fresh chicken skeletal muscles. His-YFP-Ezrin (HYE) and Capping protein were expressed in BL21EDE3* bacterial cells and purified by Ni2+ affinity and gel filtration chromatography. SLBs were prepared by fusing SUVs-DOPC: 98 mol%, Ni-NTA-DGS: 2 mol%-inside small artificial chambers glued on top of clean coverslips. Actin was polymerised by mixing dark and Atto-633-maleimide labelled g-actin in 90:10 ratio, along with capping protein, ATP, and 1xKMEH buffer to generate filaments of controlled average length. Atto-565-maleimide labelled myosin minifilaments were added along with ATP on SLB-tethered actin filaments to study its effect on actin filament organisation. The three distinct actomyosin patterns were obtained as a function of myosin density, F-actin density and F-actin length, as already reported.

### Fourier analysis of shapes

To analyze the shapes of myosin clouds we compute the Fourier spectrum of the shape boundaries [50]. The outline contour (*r*) of any given shape with a closed boundary can be expanded in terms of Fourier series as follows:

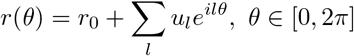

where *r*_0_ is the mean distance of any point on the boundary of the outline from the curve centroid, *u*_*l*_ is the amplitude of the *l*-th mode and *θ* is the angle made by the radius vector of such a point with the reference axis. From such an expansion we can determine the different mode amplitudes by taking a Fourier transform of *r*(*θ*) as follows:

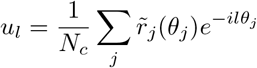

where *N*_*c*_ is the total number of points on the contour and 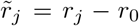, *r*_*j*_ and *θ*_*j*_ being the length of radius vector and orientation, respectively, of the *j*-th point on the contour. These modes capture specific features, for instance, *l* = 1 mode reports on the circular behavior, *l* = 2 captures the elliptic nature, *l* = 3 describes the triangular character, *l* = 4 the quadrateness and so forth.

We use the above analysis to calculate the Fourier modes of the shapes observed in our experimental images. We work with images of isolated actomyosin clusters which are subjected to a Gaussian filter in order to smoothen the boundaries. The properties of the shapes are computed using the *regionprops* function and their boundaries using *bwboundaries* function in MATLAB. We perform the Fourier analysis on the shapes within a certain size range (100-500 boundary pixels) and look at absolute values of first four harmonics of *u*_*l*_. We repeat the procedure of shape analysis on our simulated actomyosin structures. Note that the *u*_*l*_ values (heights of bars) presented in Fig. 2 and 3 are mean amplitudes from samples collected over simulated snapshots or experimental ROIs, as the case may be, and the associated errorbars represent the standard deviations across the samples.

### TIRF imaging

Imaging was performed on a Nikon Ti Eclipse microscope quipped with a TIRF module coupled to Agilent monolithic laser combiner MLC400 (laser lines: 405, 488, 561, and 653 nm; Agilent Technologies). Images were collected with a 100 *×* 1.49 NA Objective with an EMCCD camera (Evolve 512; Photometrics) yielding a pixel size of 103 nm. Multicolor, time lapse, and stream acquisition was controlled and recorded using the software *μ*Manager. Image analysis was performed in ImageJ (imagej.nih.gov/ij/).

### 3D STED nanoscopy

Stimulated emission depletion (STED microscopy) was performed on STAR-635 labelled F-actin and STAR-580 labelled myosin II on an Abberior expert line 775 nm STED system. A pulsed 775 nm depletion laser and two pulsed excitation lasers (561 nm, 640 nm) were used to image myosin and actin respectively. Spatial Light Modulator (SLM) technology on the system allowed us to tune the STED beam between lateral and axial directions to improve xy and z resolution. We characterised our fluorophores by measuring their FWHM at different 775 nm power densities. We could achieve a resolution as good as ≈ 30 nm in xy and 60 nm in z. Actomyosin asters were imaged at 50% 3D settings, with a measured resolution of 70 and 100 nm in xy, and 120 nm and 150 nm in z for actin-635 and MyosinII-580, respectively (see Supplementary Fig. S4). Signal was collected with a 100 × 1.4 NA Olympus Objective with an APD by Excelitas technologies. Images were analysed in ImageJ.

