## Supplement text and Figures for "Assemblies of F-actin and myosin-II minifilaments: steric hindrance and stratification at the membrane cortex"

### CONTENTS

S1. Simulation Methods 3
A. Coarse-grained representation 3
B. Dynamics of actin and myosin filaments 3
1. Deterministic forces on the polymer beads 4
2. Action of Myosin II motors 5
3. Turnover of myosin-II minifilaments 6
4. Stratification of Myosin II minifilaments 6
C. Units of the model 6

References 14

### S1. SIMULATION METHODS

We design our agent-based simulation using a bead-spring model for polymers with dimensions based on measured values in the *in-vitro* reconstitution experiments [1]. The main ingredients of the model are:

#### A. Coarse-grained representation

**Actin filaments:** F-actin is modelled as a linear chain polymer (Fig. S1a), whose constituent beads have a diameter of $\approx 0.14$ $\mu$m. Thus each F-actin bead represents $\approx$ 40 G-actin monomers (diameter $\approx 3.5$ nm). These dimensions are consistent with previous studies [2, 3]. Details of the magnitude of the extensional and bending spring stiffness have been discussed in the Methods. Suffice to say that our choice of bending stiffness is consistent with the measured value of the persistence length$\approx$ 16 $\mu$m [4]; this ensures that short F-actins are very rigid rods, while very long chains are semi-flexible. We use the subscript 'a' to refer to actin filaments.

**Myosin II minifilament:** Our model of the myosin-II minifilaments [5] is guided by the EM-images [1]. Each myosin-II minifilament consists of a fairly rigid backbone, attached to many motor heads arranged in helical fashion [6]. In our coarse-grained representation of a myosin-II minifilament, we construct the linear backbone by attaching two linear polymer chains of the same length side by side. This rigidly constrains the backbone structure. We then attach a side-chain bead to every alternate backbone bead. Each side-chain bead represents a collection of several motor heads. Such a construction, shown in Fig. S1a, ensures in a simple way, that the molecular length and width of myosin-II minifilaments is larger than that of the short actin filaments, consistent with observations [1, 5]. Keeping the diameter of each of the myosin beads the same as in F-actin, we find that each myosin-II head bead corresponds to 3-4 myosin-II heads. In the current model, we restrict the number of beads in a given row of the myosin backbone to 6, which makes the length of one myosin-II minifilament $l_m \approx 1$ $\mu$m. We use the subscript 'm' to refer to myosin.

#### B. Dynamics of actin and myosin filaments

We perform Brownian dynamics (BD) simulations of the model mixed-polymer system. *In the absence of activity*, this is described by the following Langevin equation for the $i^{\text{th}}$ bead,

$$m_i \frac{d^2\mathbf{r}_i}{dt^2} = \mathbf{F}_i - \zeta_i \frac{d\mathbf{r}_i}{dt} + \mathbf{F}_i^B, \tag{S1}$$

where $m_i$ is the mass of bead $i$, $\mathbf{r}_i$ is its position at time $t$, $\zeta_i$ is the friction coefficient, $\mathbf{F}_i$ is the deterministic force (i.e., forces due to interactions with other beads) and $\mathbf{F}_i^B$ is the thermal stochastic force acting on it obeying fluctuation-dissipation theorem (i.e., $\mathbf{F}_i^B = \boldsymbol{\eta}_i$, where $\boldsymbol{\eta}_i$ is an uncorrelated white noise with $\langle \boldsymbol{\eta}_i \rangle = 0$ and $\langle \eta_i^\alpha(t) \eta_j^\beta(t') \rangle = \sqrt{\frac{2k_BT}{\zeta_i}} \delta_{i,j} \delta_{\alpha,\beta} \delta(t-t')$, with $\alpha, \beta \in \{x, y, z\}$). As is it common in mesoscale simulations of biological systems, effects of inertia are ignored and we work in the overdamped limit [2, 3, 7], leading to the first-order force balance equation.

We perform the usual first-order discretization using the Euler-Maruyama algorithm leading to the update rules:

$$\mathbf{r}_i(t+\delta t) = \mathbf{r}_i(t) + \frac{1}{\zeta_i} \mathbf{F}_i \delta t + \delta r_i^B, \tag{S2}$$

where $\delta r_i^B$ is a random displacement of bead $i$ due to the Brownian force $\mathbf{F}_i^B$. It is drawn from a Gaussian distribution with zero mean and variance $2D_i\delta t$ where $D_i = k_BT/\zeta_i$, is the diffusion coefficient and $\delta t$ is the simulation time step (see Table I). Throughout our simulations we assume $\zeta_i \equiv \zeta$ for all beads. We use the relation $\zeta = 6\pi\eta\sigma$ for a bead of diameter $\sigma$, where $\eta$ is the viscosity of the medium, taken from Ref. [3].

#### 1. *Deterministic forces on the polymer beads*

The deterministic forces on each bead of the polymer, include contributions from stretching and bending of the polymer chain. The stretching energy between two connected beads $i$ and $j$ is modelled by a harmonic potential,

$$U_s(r_{ij}) = \frac{1}{2} k_s \left(r_{ij} - r_0\right)^2, \tag{S3}$$

where $k_s$ is the spring stretch constant, $r_{ij} = |\mathbf{r}_i - \mathbf{r}_j|$ is the bond length and $r_0$ is its rest length. The bending energy is described by a harmonic angular-potential,

$$U_b(\theta) = \frac{1}{2} k_b \left(\theta - \theta_0\right)^2, \tag{S4}$$

where $k_b$ is the bending stiffness, $\theta$ is the angle between two consecutive bonds meeting at bead $i$ (i.e., made by any three connected consecutive beads) and $\theta_0$ is the rest angle. We use different $k_b$ values for different types of angle which are mentioned in Table I.

Steric interactions are represented by volume exclusion due to finite size of the beads, and modelled by a truncated Lennard-Jones (LJ) potential,

$$U_r(r_{ij}) = 4\epsilon_{ij} \left[ \left( \frac{\sigma_{ij}}{r_{ij}} \right)^{12} - \left( \frac{\sigma_{ij}}{r_{ij}} \right)^{6} \right], \tag{S5}$$

where $\epsilon_{ij} \equiv \epsilon$ is the strength of the potential, $\sigma_{ij} = (\sigma_i + \sigma_j)/2$ is the effective diameter and $r_{ij} = |\mathbf{r}_i - \mathbf{r}_j|$ is the inter-bead distance. The potential is truncated at its minimum value $(-\epsilon)$ at position $r_{ij} = 2^{1/6}\sigma_{ij}$, the well-known Weeks-Chandler-Anderson form.

#### 2. Action of Myosin II motors

In our simulations, the activity of myosin involves (i) the binding of the myosin head beads minifilaments onto actin filaments, (ii) the subsequent pushing of the attached actin filaments in a directed manner with an active force and (iii) the unbinding from the actin filaments. The (un)binding of the myosin heads on actin filaments are governed by a set of stochastic mechanochemical processes such as hydrolysis of ATP, conformational changes under load and so on [8]. For simplicity, we combine these processes into two Poisson processes − binding of myosin head on F-actin with rate $k_b$ and unbinding from F-actin with rate $k_u$. To implement these rates in the simulations we assign each free (unbound) myosin head with a timescale for the next binding event, chosen from an exponential distribution with a mean equal to $t_b(\sim 1/k_b)$, the mean waiting time for binding. For each myosin head bound to F-actin, we assign another timescale for the next unbinding event, chosen again from an exponential distribution with a different mean waiting time $t_u \sim 1/k_u$. These timescales $t_b$ and $t_u$ are used as parameters in our simulations, rather than the rates. Note that these are time scales associated with individual myosin heads, not the full myosin minifilament.

In addition to these stochastic binding-unbinding kinetics, the binding of a myosin minifilament to actin filaments also involves an energetics, given by a Morse potential operating only between the heads of the myosin and beads of F-actin, which has the form,

$$U_m(r'_{ij}) = D_0 \left[ \left(1 - e^{-a(r'_{ij} - r_e)}\right)^2 - 1 \right], \tag{S6}$$

where $D_0$, $a$ and $r_e$ are the depth, width and range parameters, respectively, and $r'_{ij}$ is the instantaneous distance between the $i$-th F-actin bead and $j$-th myosin head at time $t$. This potential allows a bond of equilibriun distance $r_e$ to form between the involved beads, in this case a myosin head and a F-actin bead. The strength of the bond $D_0$, is taken to be $\approx 15\ k_BT$, which is nearly the free energy cost for hydrolysis of one molecule of ATP [9]. We do not impose any other restriction on a myosin head for attachment to an actin filament apart from the inherent steric hindrance due to finite size of the beads. The above binding energetics allows a stable binding of myosin minifilaments to actin filaments with respect to thermal fluctuations.

As long as a single myosin II head is bound to F-actin, it imparts an active force, $\mathbf{f}_a$ on the actin filament directed towards the barbed ('+') end. As a reaction, the myosin head also receives an equal opposing force, which defines the velocity of the motor head in the active state, $v_m = f_a/\zeta$. In our simulations we use $f_a$ as a parameter. We compute the average velocity of the full myosin minifilament in the state of active sliding on actin filaments (Fig. S1b). We observe that for a wide rage of forces ($2-6$ pN) the velocity varies linearly with the force. The observed velocities of the myosin minifilaments are consistent with previous studies [7].

#### *3. Turnover of myosin-II minifilaments*

When all the myosin II heads belonging to a minifilament unbind from the actin, we remove the minifilament from one part of the simulation box with a rate $k_m$, and simultaneously add another minifilament in some other part of the simulation box. This procedure introduces an effective turnover of myosin minifilaments, while keeping their number unchanged. This avoids costly memory allocations and deallocations related to the addition and removal of objects from the simulation. In the simulation the turnover is implemented with a uniform probability of turnover given by the rate $k_m \delta t$.

#### *4. Stratification of Myosin II minifilaments*

We implement stratification of the actin and myosin minifilaments, by allowing myosin minifilaments to explore the space inside a thin but finite three-dimensional (3d) rectangular box periodic along $x$ and $y$, but restricted along $z$. We restrict the actin filaments on a 2d plane with periodic boundary conditions applied along $x$ and $y$ directions, and place the plane at $z = 0$. The myosins are confined within a thin rectangular slab of thickness $3\sigma$, using a harmonic potential, $U_{slab} = 0.5 k_{slab}(z - z_0)^2$ centred at $z_0 \approx 1.5\sigma$. The myosins are also not allowed to penetrate the plane on which the actin filaments reside.

### C. Units of the model

We convert the dynamical equations to dimensionless variables by expressing all lengths in units of $\sigma$, all energies in units of the thermal energy $k_B T$ and all times in units of $\tau = \zeta\sigma^2/k_B T$. The real values of the units are displayed in Table I, which also carries information on all other parameters kept fixed throughout our simulations.

TABLE I: Parameter values in real units with dimensionless values used in simulations.

| Parameters | Real units (dimensionless values) |
|---|---|
| bead diameter, $\sigma$ | 0.14 $\mu$m (1) |
| Thermal energy, $k_BT$ | 0.0043 pN-$\mu$m (1) |
| Box length, $L_x = L_y$ | 28 $\mu$m (200) |
| Friction coefficient per bead, $\zeta$ | 0.0025 pN s $\mu$m$^{-1}$ (1) |
| Spring constant of bonds, $k_s$ | 80 pN $\mu$m$^{-1}$ (330) |
| Bending stiffness of f-actin, $k_{b,a}$ | 1.76 pN $\mu$m (400) |
| Bending stiffness of myosin backbone, $k_{b,m}^{bb}$ | 0.52 pN $\mu$m (120) |
| Bending stiffness of myosin backbone-head segments, $k_{b,m}^{bh}$ | 0.08 pN $\mu$m (20) |
| Depth of LJ potential, $\epsilon$ | 0.0043 pN-$\mu$m (1) |
| Depth of Morse potential, $D_0$ | 0.0642 pN-$\mu$m (15) |
| Width of Morse potential, $a$ | 21.43 $\mu$m$^{-1}$ (3) |
| Rest length of bonds, $r_0$ | 0.17 $\mu$m (1.2) |
| Rest length of Morse Potential, $r_e$ | 0.14 $\mu$m (1) |
| Unit of time, $\tau$ | 1.20 $\times 10^{-2}$ s (1) |
| Simulation time step, $\delta t$ | 1.80 $\times 10^{-6}$ s (0.00015) |
| Length of myosin filament, $l_m$ | 1 $\mu$m (7) |
| Strength of the harmonic slab potential, $k_{slab}$ | 48.5 pN $\mu$m$^{-1}$ (200) |
| Rate of myosin turnover, $k_m$ | 0.4 |

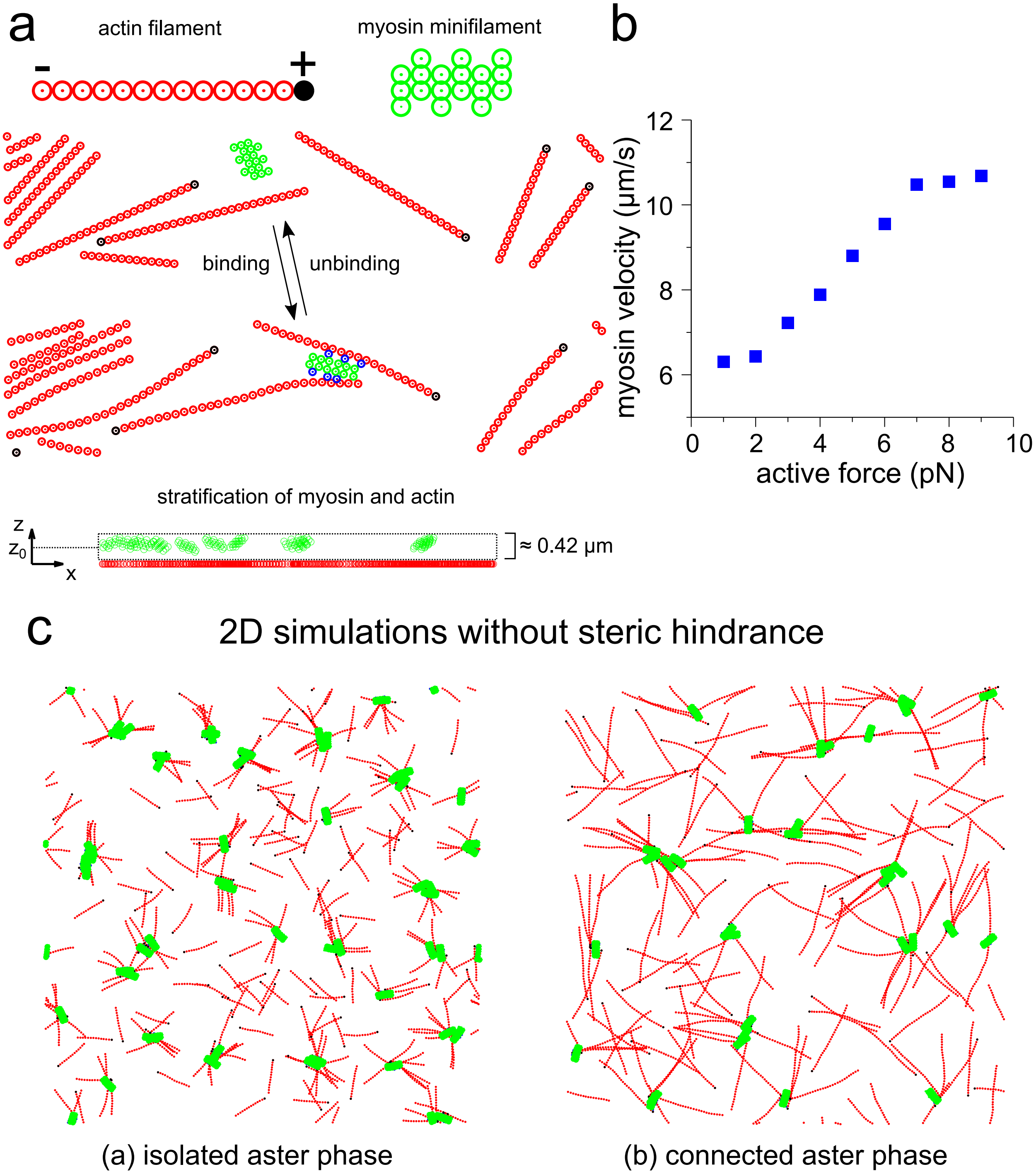


FIG. S1: (a) Description of the simulation model. (b) Velocity of myosin minifilament estimated from simulation as a function of active force. (c) Snapshots from simulations without the steric hindrance between the polymers in two dimensions (2d). The conditions are: left - $l_a = 2.32$ $\mu$m, $c_a = 1.5$ nM, $c_m/c_a = 0.2$ and right - $l_a = 4.84$ $\mu$m, $c_a = 0.8$ nM, $c_m/c_a = 0.2$. Active force is 1 pN and $k_u/k_b = 0.05$ in both cases.

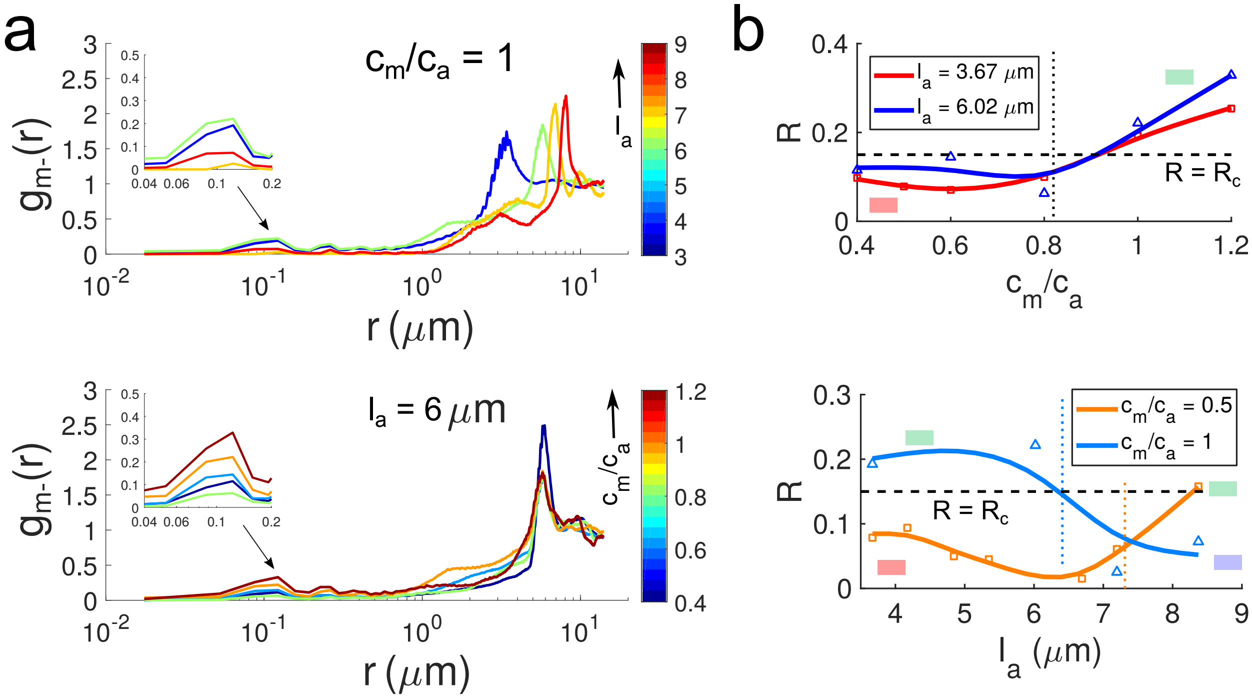


FIG. S2: Plots of the pair-correlation functions between bound myosin heads and minus ends of actin filaments, given by $g_{m-}(r)$ ('$m$' stands for myosin and '$-$' for minus-end of actin filament). We show $g_{m-}(r)$ in (a) for different $l_a$ at a fixed ratio of myosin and actin concentration ($c_m/c_a = 1$) (top), and for different values of $c_m/c_a$ for a fixed $l_a \sim 6\ \mu$m (bottom). Both the plots have insets depicting the behaviour of $g_{m-}(r)$ around $r \approx \sigma = 0.14\ \mu$m. In (b) we show few trends of the value of the first peak, our order parameter for different actomyosin cluster phases, denoted as $R \equiv g_{m-}(r \approx \sigma)$, under different conditions.

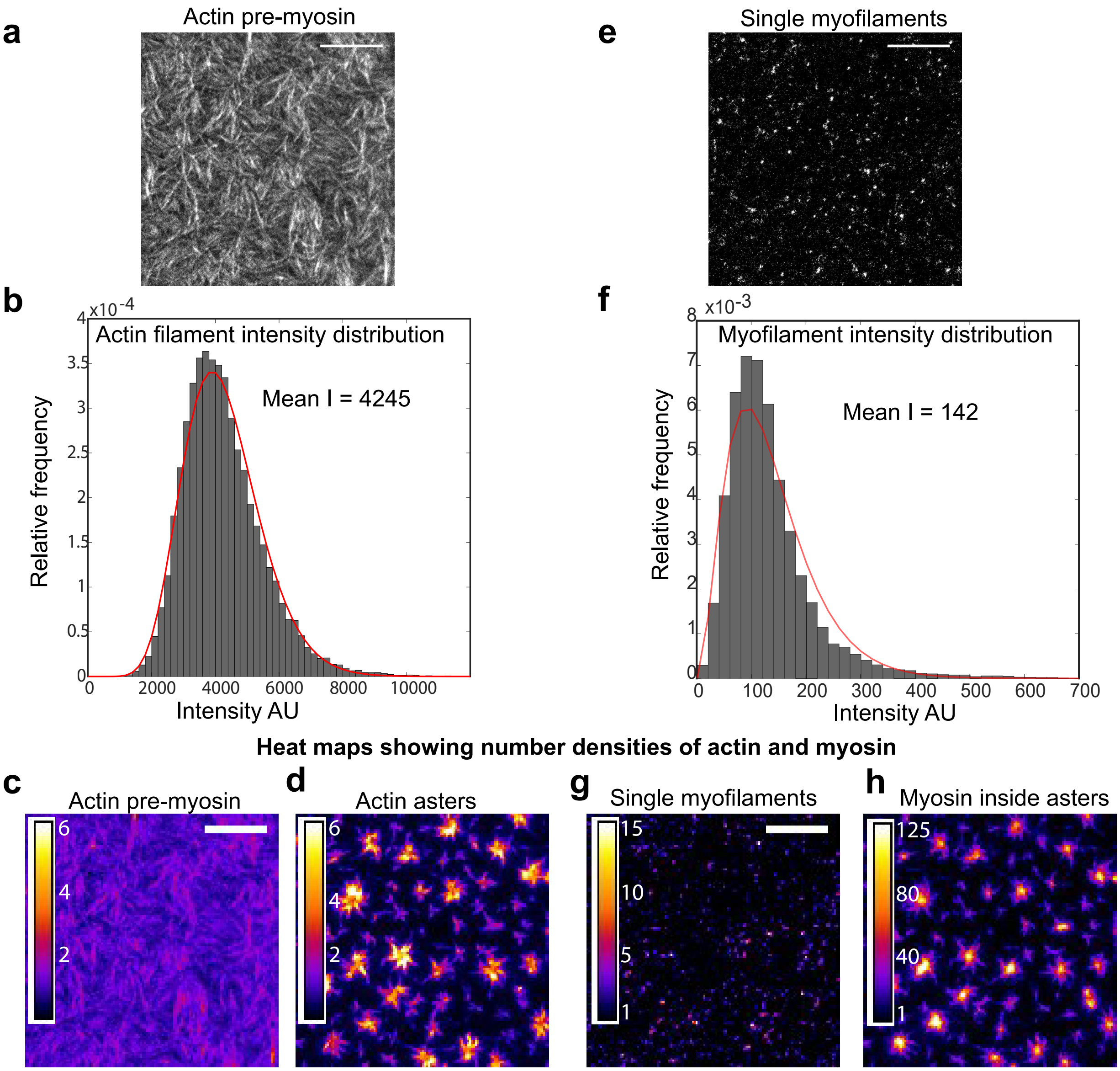


FIG. S3: Density measurements of actin and myosin in our *in-vitro* experiments. Fluorescent F-actin and myosin II filaments in a typical experiment were visualized using TIRF microscopy. (a) Snapshot of SLB bound actin filaments before myosin addition. (b) Intensity distribution of F-actin obtained from background subtracted and flat-field corrected actin images (3x3 pixels) and fitted to a gamma distribution to calculate the Mean intensity value. (c-d) All the images were then divided by the computed mean (4245 AU in this case) to generate number density maps of actin, both pre-myosin addition (c) and after aster formation (d). The calibration values on the left side of each map show the number density fold change. Radial scans of these calibrated number density maps were used to obtain circularly averaged density profile of actin asters as shown in Fig. 2g. The amplitude in y axis shows number density relative to the pre-myosin levels. (e) Snapshot of single myofilaments as they first landed on SLB bound F-actin. (f) Intensity distribution and computed Mean intensity values of single myofilaments (142 AU in this case). (g-h) Calibrated density map of single myofilaments (g) and myosin II inside asters (h). Circular density profile of myofilaments inside asters was obtained from the calibrated number density maps. Note (c) and (d) are scaled to the same intensity levels, but not (g) and (h) for better visualization

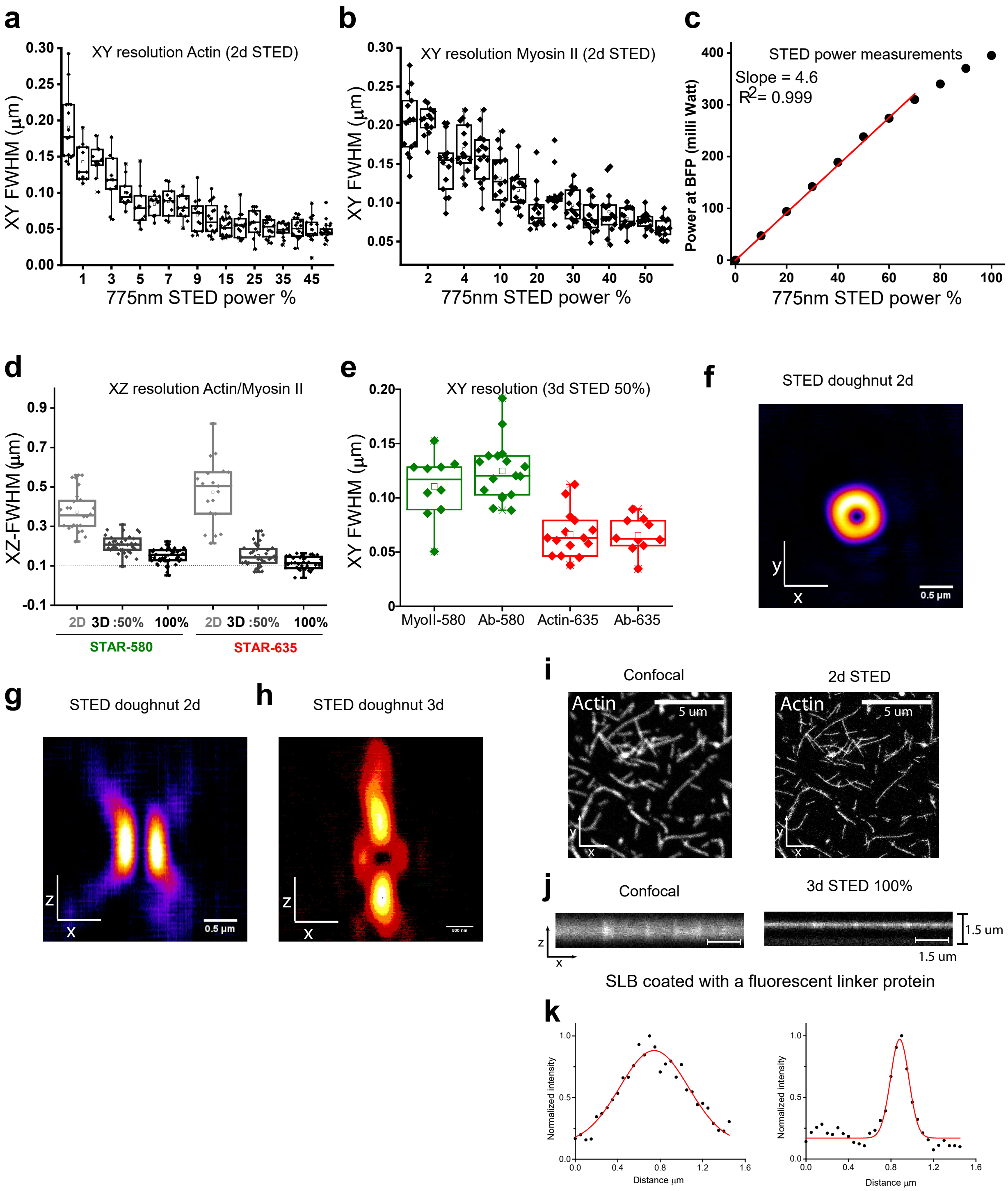


FIG. S4: Characterization of the fluorophores on Abberior 775 STED nanoscope. To probe the configurations of actin and myosin transverse to the plane of the actomyosin layer in our *in-vitro* setup, we performed 3d stimulated emission depletion (STED) microscopy on the *in-vitro* system, prepared with Star-635 labelled actin and Star-580 labelled skeletal muscle myosin II. To characterize the resolution of STED microscope, we measured the FWHM of monomeric actin and myosin molecules as well as Antibodies labelled with Star-635 and STAR-580. (a) FWHM of single g-actin molecules labeled with STAR-635 at increasing 775nm STED laser power showing improvement of xy resolution as a function of STED power. (b) FWHM of monomeric muscle myosin II (in 500 mM KCl) labeled with STAR-580 at increasing 775nm STED laser power. (c) 775nm STED power measured at the back focal plane of the objective showing linear increase till 70% laser output. With spatial light modulation, the STED doughnut shape can be modulated to improve z resolution, by steering the energy of the depletion beam from a doughnut in the xy-plane (f,g) to a shape with more depletion laser energy in the z-axis as lobes above and below the xy plane of the doughnut (h). (d) XZ FWHM of actin-635 and myosin II-580 at different STED beam shapes. (e) Box plots showing xy FWHM of STAR-635 labelled G-actin and STAR-580 labelled myosin II-580 in 3d-STED: 50% (when 50% of 775nm STED laser power was redistributed along z-axis). Note the xy resolution was better compared to the Confocal mode (a,b). We used this configuration to image our asters. (f-g) Image of a 2d STED doughnut showing 775 nm power distribution in xy (f) and xz (g). (h) Image of a 3d STED doughnut (100%) showing 775 nm power distribution in xz. (i) Comparison of actin-635 visualized under Confocal (left) and 2d STED mode (right). (j) Image of a Nickelated supported lipid bilayer (SLB) coated with a STAR-580 labeled His-tagged linker under Confocal and STED mode (3d 100%). (k) XZ intensity profile of the SLB shows improvement in z-resolution under 3d STED mode.
